# Schmallenberg virus non-structural proteins NSs and NSm are not essential for experimental infection of *Culicoides sonorensis* biting midges

**DOI:** 10.1101/2024.12.04.626779

**Authors:** Kerstin Wernike, Ana Vasic, Susanne Amler, Franziska Sick, Cristian Răileanu, Oliver Dähn, Helge Kampen, Cornelia Silaghi, Martin Beer

## Abstract

The teratogenic orthobunyavirus Schmallenberg virus (SBV) is transmitted between its mammalian hosts by *Culicoides* biting midges. The genome of circulating SBV, i.e. variants present in viraemic ruminants or insect vectors, is very stable, while variants found in malformed ruminant foetuses display a high genetic variability. It was suggested that foetal infection provides an environment that favours viral mutations that enable immune escape in the unborn, but at the costs of limiting the ability of the virus to spread further. To investigate infection and dissemination rates of different SBV variants in the insect vectors, we fed laboratory-reared *Culicoides sonorensis* with blood containing the prototype strain BH80/11-4 from a viraemic cow or strain D281/12, which was isolated from the brain of a sheep foetus and harbours multiple mutations in all three genome segments. Further, virus variants lacking NSs, NSm or both non-structural proteins were included. Six days after feeding, virus replication was found in about 2% of the midges exposed to wild-type strain BH80/11-4. The absence of the non-structural proteins had no obvious effect on the oral susceptibility to virus infection, as 2.78% of the midges fed with the NSs-deletion mutant displayed after six days viral loads higher than the respective day-0-group, 1.92% of the midges exposed to the NSm-deletion mutant and 1.55% of midges exposed to the NSs/NSm-deletion mutant. In contrast, strain D281/12 did not replicate at all in the midges, supporting the assumption that SBV variants arising in infected foetuses are unable to enter the normal insect-mammalian host cycle.

**Importance:** Biting midges are responsible for the transmission of Schmallenberg virus (SBV), a pathogen of veterinary importance that primarily infects ruminants. Although SBV has been extensively studied in the mammalian host, the virus-intrinsic factors allowing infection of, and replication in, biting midges are largely unknown. Therefore, we infected laboratory-reared *Culicoides sonorensis* midges with SBV variants by feeding them with virus-containing blood. The SBV variants differed in their genome composition, as we used the prototype wild-type strain, a strain with multiple mutations that was isolated from the brain of a malformed foetus, and recombinants lacking either NSs or NSm or both of these non-structural proteins. While the non-structural proteins had no obvious effect, the variant from the malformed foetus did not replicate at all, indicating that virus variants with characteristic genomic mutations present in foetuses lose their ability to infect the insect vector and will be excluded from the natural transmission cycle.

## Introduction

Schmallenberg virus (SBV) is an emerging pathogen that has gained significant attention in recent years due to its devastating impact on livestock, particularly cattle, sheep, and goats. First identified near the German-Dutch border region in 2011 (1), the virus spread very rapidly throughout the European ruminant population in the following years (2, 3). Acute infections are usually inapparent or associated with mild transient clinical signs such as fever, reduced milk yield or short-lasting diarrhoea (1). However, when naïve pregnant ruminants are infected during a critical phase of gestation, abortion, stillbirth or severe foetal malformation summarized under the term arthrogryposis-hydranencephaly syndrome (AHS) may be induced (4).

While the virus itself is the primary causative agent, its transmission is reliant on hematophagous insect vectors, specifically biting midges of the genus *Culicoides* (Diptera: Ceratopogonidae) (5, 6). Mosquitoes could be excluded as epidemiologically important vectors of SBV (7–9), but multiple *Culicoides* species have been implicated in virus transmission in Europe, among them *C. obsoletus*, *C. scoticus*, *C. chiopterus*, *C. dewulfi*, *C. nubeculosus* and *C. imicola* (6) (10–14). Experimental infection of laboratory-reared *Culicoides* colonies demonstrated successful dissemination of wild-type SBV in *C. sonorensis* (11).

SBV is classified as an orthobunyavirus (Simbu serogroup) within the family Peribunyaviridae (15). Like a typical bunyavirus, SBV has a negative-sense RNA genome consisting of three segments. The large (L) segment encodes the viral RNA-dependent RNA polymerase (RdRp), which is crucial for transcription and replication of viral RNA (4, 16). The small (S) segment encodes the nucleocapsid (N) protein and the non-structural protein NSs. The N protein is essential for packaging the viral RNA and protecting it from host immune defence. In addition, it plays a role in transcription and replication of the viral genome (16). NSs is a major virulence factor in the mammalian hosts, as it counteracts the shutoff of host cell protein synthesis and the induction of interferon (IFN). For the insect vector, the role of the viral NSs protein was not yet elucidated in detail, but experimental infection studies of mosquito cell lines and *Ae. aegypti* mosquitoes with Bunyamwera virus (BUNV), another member of the genus *Orthobunyavirus*, suggest that the NSs is required for efficient replication and dissemination in mosquitoes (17).

The medium (M) segment encodes the two glycoproteins Gn and Gc and the non-structural protein NSm. The function of NSm in the viral life cycle is still largely unknown. For SBV, it has been demonstrated that NSm is dispensable for virus replication in mammalian cells and cattle (18). Further, an NSm-deletion mutant of Oropouche virus (OROV), another orthobunyavirus closely related to SBV, displayed similar *in vitro* growth characteristics as the wild-type virus in mammalian and mosquito cells (19). The glycoproteins Gn and Gc are involved in viral entry by binding to host cell receptors. Within the Gc coding region of the M-segment, a region of high genetic diversity was identified (“Gc 234” or “Gc head”) (20, 21). Interestingly, an increase in genetic variation can only be observed in viruses from malformed foetuses, while the overall stability of virus variants found in viraemic animals or infected insects is very high (3, 22–24). It has been proposed that the mutations in the “hot spot” within the M-segment leads to efficient immune escape from neutralizing antibodies in infected foetuses. However, these mutant virus strains from foetuses are considered to be dead-end variants that are not able to spread further, because they have never been detected circulating, i.e. in the insect vectors or in viraemic mammalians (25). In addition, cell-culture based studies suggest that highly variable virus variants from malformed foetuses have an impaired *in vitro* growth in *Culicoides* cell lines (26). However, definite evidence about their replication characteristics in insects, e.g. from experimental infection of biting midges, is missing.

Vector competence studies play a crucial role in understanding the interaction between pathogens and their insect vectors, whereat vector competence refers to the ability of an insect vector to acquire, maintain, and biologically transmit a specific pathogen. To achieve transmission between susceptible mammalian hosts by insects, an arbovirus needs to be able to both replicate and disseminate within the vector (27). In addition, experimental infections enable detailed investigations of the role of specific proteins or protein domains of the pathogens within the insects.

In the present study, we exposed *C. sonorensis* biting midges to SBV by oral ingestion in order to investigate the colonization and replication properties of different virus variants, among them a strain isolated from a malformed newborn and viruses lacking the NSs and/or NSm protein.

## Materials and methods

### Virus variants used for infection

The SBV strain BH80/11-4 was isolated from a viraemic cow in 2011 using baby hamster kidney (BHK) and *C. sonorensis* (KC) cells (1). The virus strain SBV D281/12 was isolated from the brain of a malformed foetus on BHK cells (26). The full-length sequences of the S, M and L segments are available at NCBI GenBank (strain BH80/11-4: accession numbers HE649914 (S), HE649913 (M) and HE649912 (L); strain D281/12: accession numbers PP626413 (S), PP616750 (M) and PP626412 (L)). SBV D281/12 harbours multiple point mutations in all three genome segments and a large genomic deletion in the M-segment (S1 Supplementary Figure). Previous experiments have shown that BH80/11-4 replicates well in KC cells, while D281/12 has an impaired *in vitro* growth in this insect cell line (26).

In addition, recombinant SBV (rSBV) was used for the biting midge infection experiment. Recombinant viruses based on the sequence of the strain BH80/11-4, and either unmodified rSBV BH80/11-4 or virus variants lacking the NSs (rSBV BH80/11-4 dNSs), NSm (rSBV BH80/11-4 dNSm) or both non-structural proteins (rSBV BH80/11-4 dNSs/dNSm) were generated in previous studies (18, 28). The genomic alterations in comparison to the wild-type virus are shown in S1 Supplementary Figure.

All virus variants were propagated on BHK cells (RIE164, Collection of Cell Lines in Veterinary Medicine (CCLV), Friedrich-Loeffler-Institut, Greifswald-Insel Riems, Germany). For oral infection of the midges, virus preparations containing 10^6 50 % tissue culture infective dose per ml (TCID_50_/ml) were produced. The titres were confirmed by back-titrations of leftovers of the inocula on BHK cells subsequent to feeding.

### Rearing of Culicoides sonorensis

A laboratory colony of *Culicoides sonorensis* was reared in the BSL2 insectary of the Friedrich-Loeffler-Institut, Greifswald-Insel Riems, Germany, under standardized conditions (25±1 °C, 70 % relative humidity, 12:12-hour light cycle including one hour each dusk and dawn). The rearing cycle included egg collection, development of larvae, collection of pupae and production of adult midges. Larvae were kept in 30x41x10 cm^3^ plastic containers with an elongated sponge-made central island, and a constant water flow produced by an electrical rotating device. Larvae were fed with ground fish food (TetraMin flakes grounded, Tetra, Melle) every two days. Pupae that concentrated on the sponge island were collected using addition of water and mechanical collection from the surface to a 500 ml jar. The vertical cage system for development and maintenance of adult midges consisted of 50 ml conical tubes filled with water and a carton cage with a hole fitting the conical tube at the bottom and a net on the top. Pupae were transferred onto the filter paper on the top of the tube, allowing humidity to pass from the water in the tube. For colony maintenance, adult *C. sonorensis* were fed on 37 °C warm ovine blood (provided by Friedrich-Loeffler-Institut) using a Hemotek membrane feeding system (Hemotek, Blackburn, UK) three times a week, and produced eggs were collected on the filter paper which was placed in the water of a larval rearing container.

### Experimental design and sample processing

Three day-old mixed male and female adult *C. sonorensis* were kept at 27 °C on a 12:12 hour light cycle with 10 % fructose sugar solution provided ad libitum prior to exposure to SBV. Biting midges were offered ovine (heparin) blood mixed 1:1 with SBV in cell culture medium. The ovine blood had been confirmed to be negative for SBV genome using real-time RT-PCR (RT-qPCR) (29) and for anti-SBV antibodies by ELISA (ID Screen® Schmallenberg virus Competition Multi-species, Innovative Diagnostics, Grabels, France) before merging it with the cell culture supernatant containing the SBV variants. The midges were allowed to feed for 30 min on the blood prewarmed to 37 °C using a Hemotek membrane feeding system. Afterwards, obviously unfed individuals were sorted out using bare eyes under short-term anaesthesia (CO_2_) on a cooling plate. Only clearly engorged midges were kept for the course of the experiment.

In the first infection experiment, the wild-type viruses BH80/11-4 and D281/12 were compared to each other regarding their ability to replicate in the insects. A total of 938 midges were fed with a blood preparation containing SBV BH80/11-4 and 530 midges with blood containing SBV D281/12. In addition, rSBV BH80/11-4 was included (208 midges) in order to demonstrate the equality of the recombinant to the wild-type virus. As negative controls, 440 midges were fed with SBV-free ovine blood. Subsamples of the blood-fed midges were processed immediately after feeding, others after 4, 6 or 8 days of incubation at 27 °C (Figure 1). The total number of midges per day and group, which were tested in toto, is indicated in Figure 2. In addition, 252 midges of the BH80/11-4 group were dissected and heads and bodies were analysed separately.

**Figure 1:**
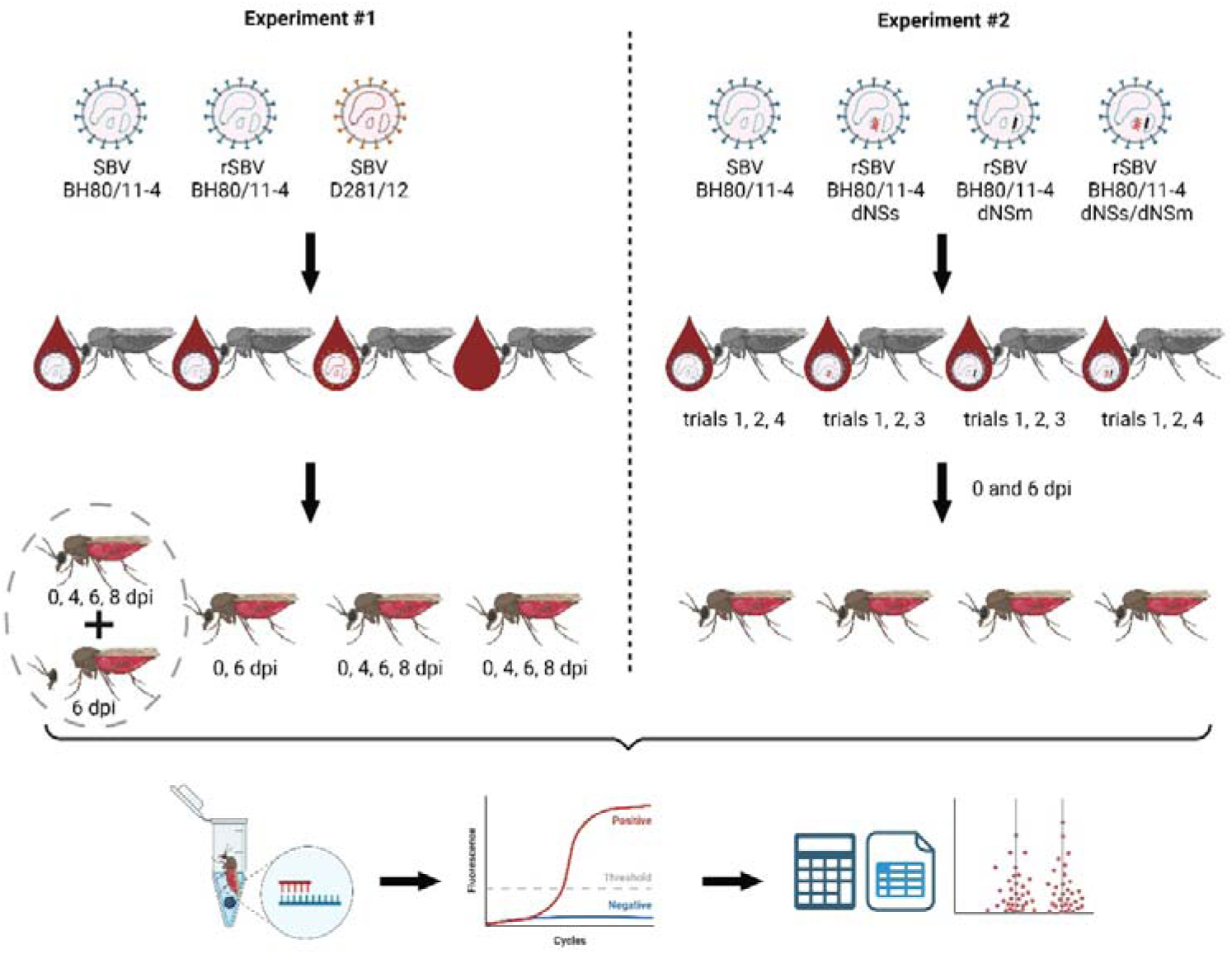
Experimental design. In the first experiment, midges were infected with blood containing SBV wild-type strains BH80/11-4 or D281/12 or recombinant BH80/11-4. A control group was fed with SBV-free ovine blood. For the second experiment, the recombinant virus variants rSBV BH80/11-4, rSBV BH80/11-4 dNSs, rSBV BH80/11-4 dNSm and rSBV BH80/11-4 dNSs/dNSm were used for infection of midges by blood meal. At the indicated days post infection (dpi), viral RNA was extracted from each midge individually and analysed by an SBV-specific RT-qPCR. For a subset of midges infected with wild-type strain BH80/11-4 body and head were processed separately. The generated data were analysed as describe in the materials and methods section. Created with BioRender.com

**Figure 2:**
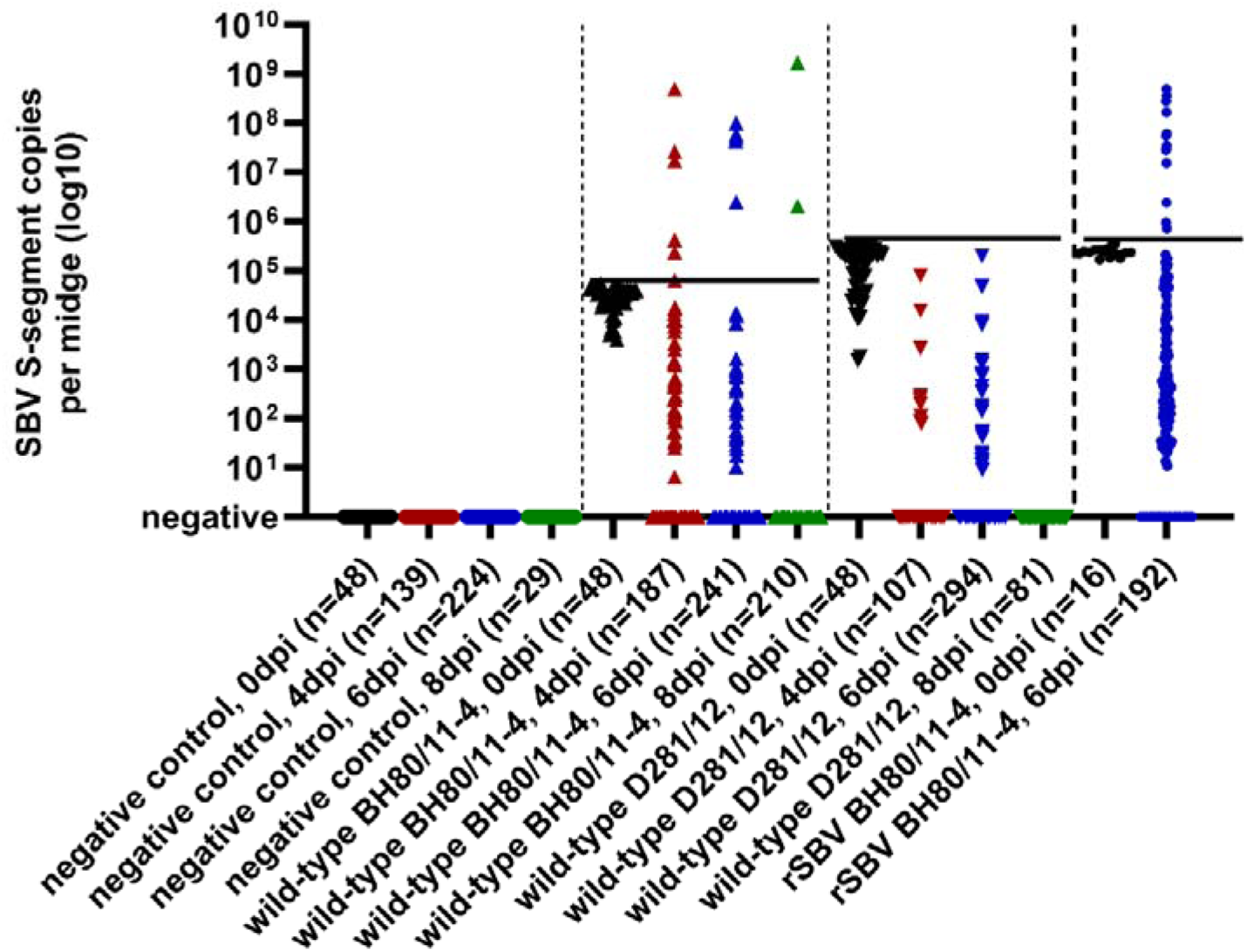
RT-qPCR results of midges experimentally infected with the wild-type SBV strain BH80/11-4, the same strain produced by a reverse genetics system (recombinant SBV, rSBV), or by the virus variant SBV D281/12, and midges fed with virus-freed blood. Individual midges were tested either immediately after ingesting the SBV-containing blood meal (0 dpi, black) or four days (4 dpi, red), six days (6 dpi, blue) or eight days (8 dpi, green) post infection. Horizontal black lines indicate the highest SBV S-segment copy number measured in any of the midges of the respective group immediately after ingesting the SBV-containing blood meal.

In the second experiment, rSBV and the deletion mutants rSBV BH80/11-4 dNSs, rSBV BH80/11-4 dNSm and rSBV BH80/11-4 dNSs/dNSm were fed to the midges (Figure 1). Because of the large numbers of individual midges included in this comparison, the experiment was split into four trials. The number of midges per trial and virus variant is given in Fig 3. Based on the results of the first experiment, only two time points were selected for the PCR analysis of the midges, namely 0 and 6 days post infection (dpi).

**Figure 3:**
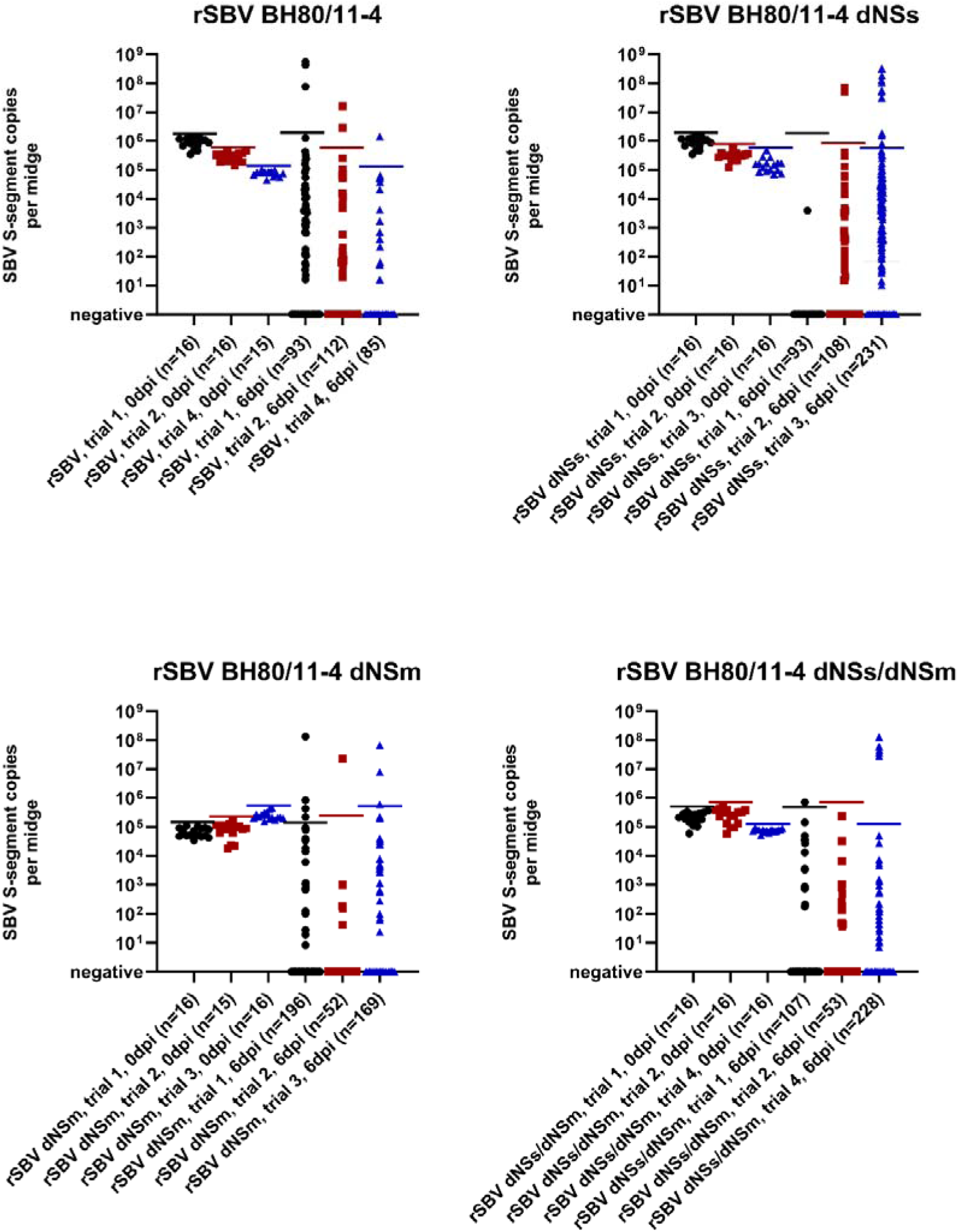
RT-qPCR results of midges experimentally infected with rSBV BH80/11-4 or recombinant mutants lacking the non-structural protein NSs (rSBV BH80/11-4 dNSs), NSm (rSBV BH80/11-4 dNSm) or both of these proteins (rSBV BH80/11-4 dNSs/dNSm). Midges were tested for SBV genome either immediately after blood meal or 6 days post infection (6 dpi). Horizontal lines indicate the highest SBV S-segment copy number measured in any of the midges of the respective group immediately after ingesting the SBV-containing blood meal. The experiment was performed in 4 subsequent trials, the trial number is given in the label of the x-axis. In the individual figure panels, results for midges from the identical trial within this experiment are indicated by the same colour.

To be used for RT-qPCR analyses, total RNA was extracted from each midge individually. Insects were homogenized in 200 µl Schneider’s medium or phosphate-buffered saline (PBS) using one 5 mm stainless steel ball per midge and a TissueLyser (Qiagen, Hilden, Germany) agitating for one minute at 30 Hz. Total nucleic acid was extracted from 100 μl of the homogenates using the King Fisher 96 Flex (Thermo Scientific, Braunschweig, Germany) in combination with the NucleoMag VET kit (Macherey-Nagel, Düren, Germany) according to the manufacturer’s instructions. Subsequently, the extracts were analysed by an S-segment based RT-qPCR (29) with an external standard, which was used to calculate the number of SBV genome copies per midge. For a subset of midges infected with wild-type BH80/11-4 (n=252), heads and bodies were processed separately.

### Statistical analysis

The primary endpoint for all exploratory analyses was the occurrence of virus replication at three independent time points 4, 6 or 8 days after feeding, defined as present when a predetermined threshold value of genome copy numbers for the respective day-0-group is exceeded. Accordingly, it was investigated whether the proportion of midges with higher values in one experimental virus variant group was significantly greater than the null proportion, which means that virus replication has taken place in at least one midge. In order to detect virus replication, the exact one-sided binomial test for one proportion was used to determine if the observed proportion met the expected benchmark of at least one midge with viral loads higher than the day-0-group. In a second step, two-sided Fisher exact test was used to test for differences in the proportions of midges with virus replication between wild-type SBV strain BH80/11-4 and the different virus variant groups. P-values less than 0.05 indicate statistical significance. Statistical analyses were carried out in the open-source software environment R (30) using the package binom (31).

## Results

### Experiment #1: Wild-type SBV from a viraemic cow, but not a virus variant from a malformed lamb, replicates in C. sonorensis

Exposure of midges to the prototype wild-type SBV strain BH80/11-4 by ingestion of an infective blood meal led to PCR positivity in all engorged insects immediately after feeding. The highest genome copy number in an individual of this group was used as cut-off for the intake background to assess virus replication during the following days in the other midges fed with the same virus preparation. At 4 dpi, 48 out of 187 midges tested positive by RT-qPCR and six of them (6/187, 3.21 %) displayed viral loads higher than the day-0-group, indicating virus replication. At 6 dpi, 31 out of 241 midges gave positive results, five (5/241, 2.07 %) with values above the cut-off. At 8 dpi, two midges tested positive and both of them harboured viral loads considerably above that of the day-0-group (2/210, 0.95 %) (Figure 2). Infection of midges with rSBV BH80/11-4 led to similar results, the 16 engorged individuals that were tested immediately after feeding scored positive by RT-qPCR. Of the 192 midges analysed at 6 dpi, 12 (6.25 %) gave positive results with genome copy numbers higher than the respective day-0-group. These proportions were slightly different compared to wild-type BH80/11-4 at 6 dpi (2.07 % vs. 6.25 %, p = 0.043).

To further assess vector competence for wild-type BH80/11-4, head and body homogenates of 252 midges collected at 6 dpi were analysed individually. Both, head and body, tested negative in 223 (88.49 %) cases. A total of 29 midges gave positive results in the SBV RT-qPCR, in 12 cases only the body scored positive, in eight cases only the head and in nine cases both. In six midges (2.38 %), the viral load exceeded that of 0 dpi of the wild-type BH80/11-4 infected group (Figure 4).

**Figure 4:**
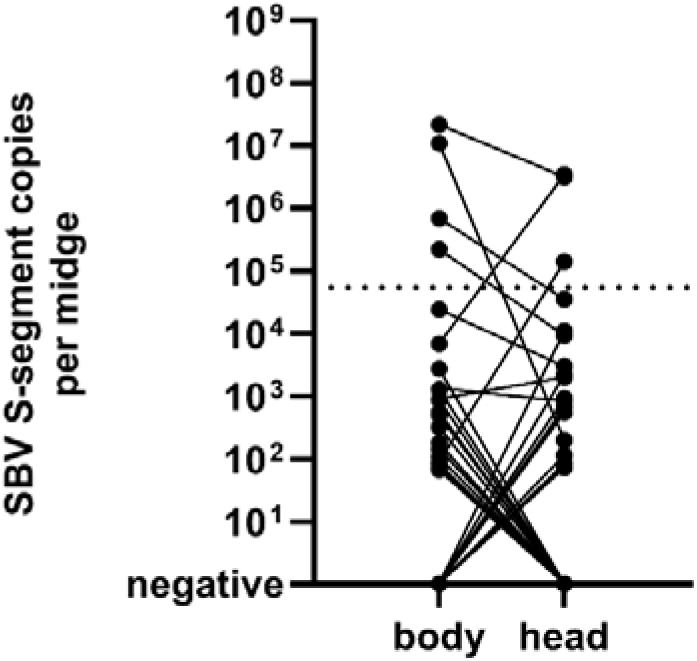
Comparison of SBV S-segment copies measured in bodies and heads of experimentally infected midges. The results of the body and of the head of an individual midge are connected by a black line.

In contrast to wild-type BH80/11-4 and rSBV BH80/11-4, the virus strain SBV D281/12 did not replicate in midges. Though all engorged midges tested positive immediately after ingesting the infectious blood meal, all individuals analysed in the following days gave either negative RT-qPCR results or were tested positive with genome copy numbers below the cut-off for this infection group (= highest genome copy number in an individual immediately after feeding with D281/12-containing blood). This observed difference in proportions compared to wild-type BH80/11-4 was statistically significant at 6 dpi (0/294, 0 % vs. 5/241, 2.07 %, p = 0.018) (Figure 2).

### Experiment #2: SBV non-structural proteins NSs and NSm did not have an apparent effect on virus replication characteristics in biting midges

To investigate the role of SBV’s non-structural proteins for replication in midges, the insects were exposed to virus variants lacking NSs, NSm or both proteins. Again, the highest genome copy number in an individual of a given group and trial was used as cut-off for the day-6-values of this respective group to assess virus replication.

Like in the previous experiment, ingestion of an infective blood meal led to PCR positivity in all engorged insects immediately after feeding, independent of the virus variant (Figure 3).

Six days after exposure to unmodified rSBV BH80/11-4, 2.07 % of the midges gave positive results with genome copy numbers higher than the respective day-0-group (3/93 in trial 1, 2/112 in trial 2 and 1/85 in trial 4) (Figure 3).

Ingestion of SBV variants lacking one or both non-structural proteins also led to virus replication in a subset of insects. Six days after exposure to rSBV BH80/11-4 dNSs, 2.78 % of the midges displayed viral loads higher than the day-0-groups (0/93 in trial 1, 2/108 in trial 2 and 10/231 in trial 3). Six days after ingestion of blood containing the virus variant rSBV BH80/11-4 dNSm, 1.92 % of the midges gave positive results with genome copy numbers higher than the day-0-groups (4/196 in trial 1, 1/52 in trial 2 and 3/169 in trial 3). rSBV BH80/11-4 dNSs/dNSm, the virus variant lacking both non-structural proteins, was detected with viral loads higher than the respective day-0-groups in 1.55 % of the engorged midges at 6 dpi (1/107 in trial 1, 0/53 in trial 2 and 5/228 in trial 4). There were no significant differences in the proportions of virus replication between the rSBV BH80/11-4 control group and the respective virus variant in any of these three trials (Figure 3).

## Discussion

Experimental infection studies are essential to elucidate vector competence and vector-pathogen interactions, including the efficiency of virus replication and dissemination under given environmental conditions. Here, we investigated the replication characteristics of different SBV variants in *Culicoides* midges. As infection route, ingestion of an infective blood meal was selected. An alternative technique would have been intrathoracic injection of a virus suspension into the insects, which might be considered more reliable regarding virus uptake by every individual and is appropriate to examine vector-pathogen interaction. However, for vector competence studies the value of this method is limited, as it circumvents the host’s midgut barriers (32).

Another challenge in vector studies, be it experimental infection or field studies, is the interpretation of real-time PCR values (11). During the initial phase of infection, low subtransmissible viral loads in the vector are common and, especially when using high virus doses for experimental infection, viral RNA detected in the insect could originate from the original blood meal. Usually, blood digestion is complete after a few days (33, 34). Hence, viral RNA detected after that period indicates successful infection, while the measured background level confirms virus uptake via the blood meal. When digestion is not yet completed, comparison with the quantities of virus in the original blood meal can be used to demonstrate replication in the putative vector. Another option, which has been applied in this study, is to compare the viral loads to a group of midges that ingested the identical virus preparation and was tested immediately after ingestion. Higher SBV genome copy numbers at later time points clearly indicate virus replication within the insect.

As *Culicoides* species we chose *C. sonorensis*. As opposed to the European *Culicoides* species considered important SBV vectors in nature (35), this species can be reared in the laboratory and is widely used as a laboratory model biting midge. Successful dissemination of wild-type SBV in *C. sonorensis* was reported previously (11), which could be confirmed in this study for a wild-type strain isolated from viraemic cattle blood. Similarly, virus replication was detected in some midges inoculated with recombinant SBV lacking either NSs, NSm or both non-structural proteins. In mammals, the NSs protein of orthobunyaviruses has been exhaustively investigated and demonstrated to be a major virulence determinant, as it antagonizes the antiviral response by blocking the type I interferon production (16). In ruminants, a lack of the NSs protein or a combined NSs/NSm deletion leads to a replication deficit; the deletion mutants do not induce detectable virus replication or clinical disease in cattle anymore (18). In addition, immunization with the avirulent double deletion mutant fully protects from virulent wild-type virus challenge infection. Hence, the mutant viruses were proposed as candidate vaccines (18). In this context, the question arose whether the deletion variant viruses can be transmitted by the insect vectors, which could be an additional safety aspect if no transmission occurs. From *in vitro* studies using insect cell lines and different bunyaviruses, contradictory results are available. While experimental infection studies of mosquito cell lines with BUNV suggested that a lack of NSs seemed to delay the progress of infection in *Ae. aegypti* Ae cells, NSs was non-essential for growth in *Ae. albopictus*-derived C6/36 and C7-10 cells (17). Further, no specific function was found for the NSs protein of La Crosse orthobunyavirus (LACV) in mosquito cells (36), and OROV lacking NSs grew to similar levels on *Ae. albopictus* and *Ae. aegypti* cells as the wild-type virus (19). *In vivo*, BUNV-NSs seems to be required for efficient replication and dissemination, as fewer *Ae. aegypti* mosquitoes were infected with the NSs-deletion mutant, and the wild-type virus disseminated to salivary glands more efficiently (17). However, for the phlebovirus (family *Bunyaviridae*) Rift Valley fever virus (RVFV), no differences in infection or dissemination rates were found for a wild-type variant and an NSs-deleted virus in *Ae. aegypti* (37). Data from experimental infections of midges instead of mosquitoes with bunyaviruses other than SBV lacking the NSs protein are unfortunately not available. Hence, the aforementioned experimental infections of mosquitoes are the only studies usable for comparison with our results, but when comparing studies in mosquitoes and in midges one needs to keep in mind the differences in the biology of both. As an example, in midges, in contrast to mosquitoes, no salivary gland infection barrier is believed to exist (27, 38), simplifying the assessment of the dissemination.

Here, we could show that the NSs protein of SBV has no obvious effect on virus replication in *C. sonorensis*. Similarly, a deletion of the NSm protein did not prevent infection and replication, which is in line with cell culture-based results using the closely related Simbu serogroup virus OROV. On *Ae. albopictus* and *Ae. aegypti* cells, OROV with and without NSm displayed similar growth characteristics, suggesting that NSm is dispensable for virus replication (19). In contrast, deletion of NSm from RVFV nearly abolished the virus’ ability to replicate in *Ae. aegypti* mosquitoes (37). The molecular basis for the differences is still largely unknown and should be investigated in future studies in order to design new approaches to control bunyaviruses. Furthermore, to ultimately prove vector competence for specific virus variants, transmission experiments from the vector species to mammalian hosts are required, as has been performed with *Culicoides* spp. and, for example, the wild-type variants of the Simbu serogroup viruses OROV and Akabane virus (AKAV) (39, 40).

In contrast to the SBV variants lacking the non-structural proteins, the virus variant D281/12 from a malformed foetus did not replicate in midges. This virus strain harbours multiple mutations in all three genome segments, with a large deletion in the M-segment being the most prominent mutation. In infected ruminant foetuses, mutations within the Gc head domain are supposed to lead to immune escape from neutralizing antibodies (24, 25), but the role in insects is largely unknown. However, to focus on the large, eye-catching deletion could even be misleading, as vector competence might be related to a single point mutation. For the midge-transmitted reovirus bluetongue virus, vector competence is strongly affected by a small deletion or point mutations (41), and a single amino acid mutation in the genome of the mosquito-transmitted chikungunya virus led to a significant change in vector specificity resulting in a major disease outbreak with thousands of human cases (42). For SBV, *in vitro* studies using *C. sonorensis* cells suggest that a point mutation in the S-segment is responsible for loss of viral fitness in the insect hosts (26), which should be confirmed in the future by experimental infection of midges with chimeric viruses that contain individual mutations of strain D281/12 in the backbone of the wild-type virus. But whatever mutation it may be due to, the missing replication of the strain D281/12 in *Culicoides* biting midges supports the assumption that SBV variants with characteristic genomic mutations arising in infected ruminant foetuses are not able to enter the usual insect-mammalian host cycle and, as a consequence, cannot spread further.

## Acknowledgements

*Culicoides sonorensis* were originally developed and supplied by The Pirbright Institute under BBSRC project code: BBS/E/I/00007039. We thank Bianka Hillmann, Gabriela Adam and Oliver Tauchmann for excellent technical assistance.

This work was supported by the German Federal Ministry of Food and Agriculture (BMEL) through the Federal Office for Agriculture and Food (BLE) under Grants 281B101816 (to HK and MB) and 28N207601 (to HK and MB). The funders had no role in study design, data collection and analysis, decision to publish, or preparation of the manuscript.

## Author Contributions

Conceptualization: Kerstin Wernike, Helge Kampen, Cornelia Silaghi, Martin Beer.

Formal analysis: Kerstin Wernike, Susanne Amler.

Investigation: Kerstin Wernike, Ana Vasic, Susanne Amler, Franziska Sick, Cristian Răileanu, Oliver Dähn.

Methodology: Kerstin Wernike, Ana Vasic, Susanne Amler, Cornelia Silaghi, Martin Beer.

Supervision: Kerstin Wernike, Helge Kampen, Cornelia Silaghi, Martin Beer.

Visualization: Kerstin Wernike.

Writing – original draft: Kerstin Wernike. Writing – review & editing: All authors.

---

**Supplementary Figure S1:**
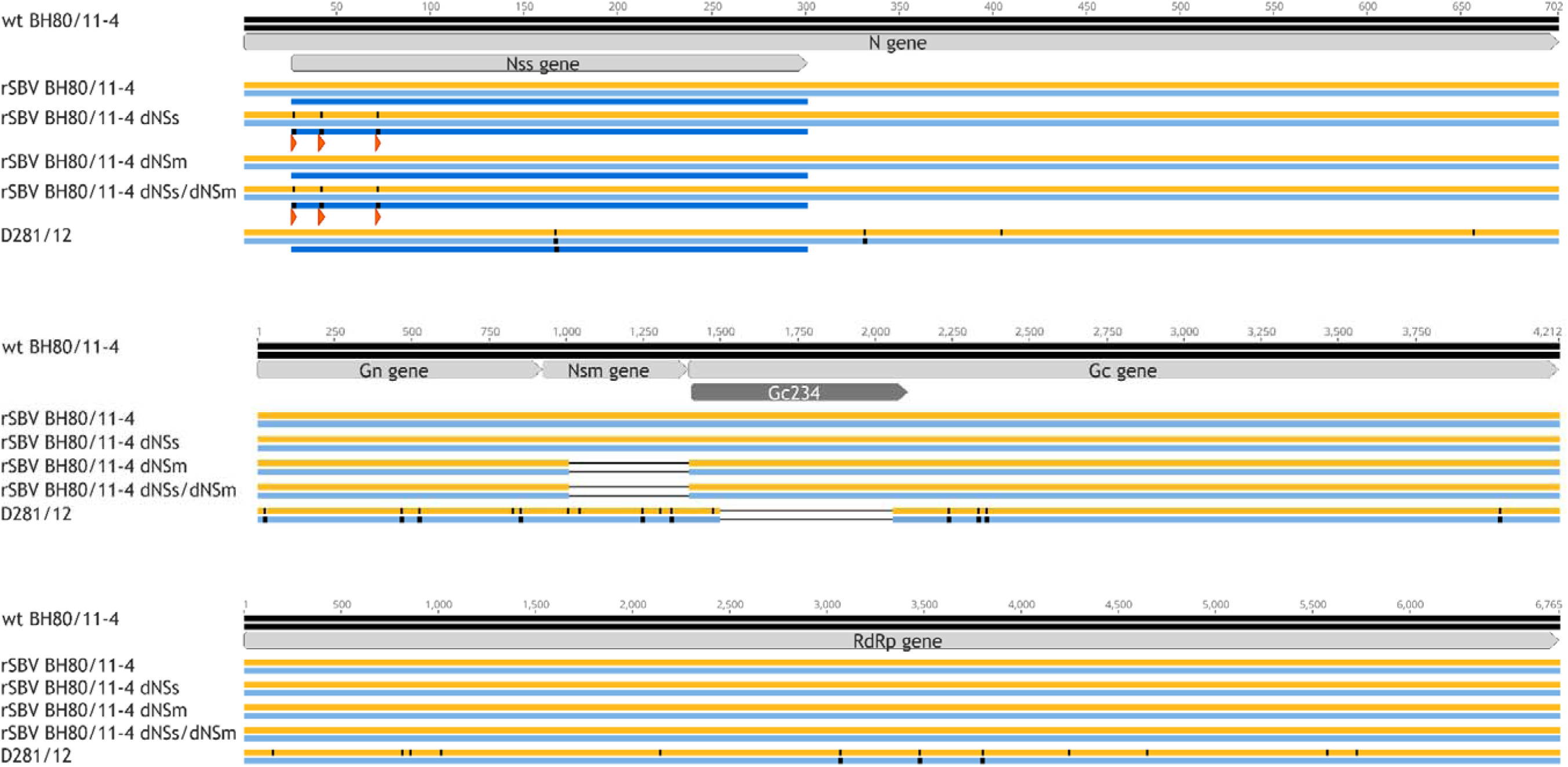
Comparison of the protein-coding regions of the S-segment (upper panel), M-segment (middle) and L-segment (lower panel) sequences of the SBV strains used in this study. The sequences of the wild-type (wt) strain SBV BH80/11-4 are used as reference. Nucleotide (orange bar alongside the name of each SBV variant) and amino acid (blue bars) substitutions are highlighted as vertical black lines. For the S-segment, the amino acid sequences are given separately for the N-protein (light blue) and for the non-structural protein NSs (dark blue), which is encoded in an alternative overlapping reading frame. Red triangles indicate mutations within the three potential translation start codons of the NSs-encoding sequence that abrogate the expression of the NSs-protein.

